# Refining Convergent Rate Analysis with Topology in Mammalian Longevity and Marine Transitions

**DOI:** 10.1101/2021.03.06.434197

**Authors:** Stephen Treaster, Jacob M. Daane, Matthew P. Harris

## Abstract

The quest to map the genetic foundations of phenotypes has been empowered by the modern diversity, quality, and availability of genomic resources. Despite these expanding resources, the abundance of variation within lineages makes the association of genetic change to specific phenotypes improbable. Drawing such connections requires an *a priori* means of isolating the associated changes from background genomic variation. Evolution may provide these means via convergence; i.e., the shared variation that may result from replicate evolutionary experiments across independent trait occurrences. To leverage these opportunities, we developed *TRACCER: Topologically Ranked Analysis of Convergence via Comparative Evolutionary Rates*. As compared to current methods, this software empowers rate convergence analysis by factoring in topological relationships, because variation between phylogenetically proximate trait changes is more likely to be facilitating the trait. Pairwise comparisons are performed not with singular branches, but in reference to their most recent common ancestors. This ensures that comparisons represent identical genetic contexts and timeframes while obviating the problematic requirement of assigning ancestral states. We applied TRACCER to two case studies: marine mammal transitions, an unambiguous trait which has independently evolved three times, as well as the evolution of mammalian longevity, a less delineated trait but with more instances to compare. TRACCER, by factoring in topology, identifies highly significant, convergent genetic signals in these test cases, with important incongruities and statistical resolution when compared to existing convergence approaches. These improvements in sensitivity and specificity generate refined targets for downstream analysis of convergent evolution and identification of genotype-phenotype relationships.

## Introduction

When challenged with similar selective pressures, independent lineages may converge on similar adaptations to those challenges (Losos, 2011). A myriad of complex mutations can be sufficient to produce such adaptive traits, but the stochastic nature of evolution begets that such adaptive traits will tend to manifest parsimoniously, particularly given the genetic constraints and largely similar toolkits of life (Rosenblum et al., 2014). Given these developmental and physiological constraints in organisms, the convergent evolution of traits will tend towards similar molecular mechanisms, providing a valuable map for phenotype to genotype inferences, recently coined “forward genomics” (Currie, 2013; Hiller et al., 2012). However, while the parsimonious path to the same adaptive phenotype may tread upon the same genetic elements or pathways, the specific molecular changes may evolve in unique ways. Given a convergent trait, the molecular similarities will be shaped both by the genetic context and the precision required; there are countless ways to disrupt a pathway or knockout a gene, but there may only be a handful to specifically compromise a hydrophobic pocket, and only one to create a disulfide bridge between two domains. A variety of changes could modulate cellular activity in comparable ways and ultimately manifest as convergent phenotypes of interest. Analytical tools with a more general measure are needed to capture these disparate genetic signatures acting on the same system in order to make meaningful comparisons between species.

### Relative Evolutionary Rates and Convergence

The assessment of relative evolutionary rates (RERs) has arisen as a powerful and flexible way to distill and compare the variable signatures acting on a system to identify convergence. RERs describe the degree of change in a genomic region as compared to the background rate across that genome. Not all parts of a genome evolve at the same pace, so these relative rates can vary widely. While mutations are randomly distributed, the finely tuned core pathways will undergo strong purifying selection, leaving them relatively unchanged, sometimes for hundreds of millions of years (Katzman et al., 2007). These loci will reliably yield low RERs. Regions in the wilderness of “nonfunctional junk” DNA can accumulate mutations freely, without selective pressure, yielding high RERs. Duplicated and pseudogenized regions, also freed from restriction, can evolve rapidly and neofunctionalize to facilitate adaptation or go to drift, and may also yield high RERs. These accelerations and constraints of change on a locus can be powerfully informative as to the selective pressures acting on it (Echave et al., 2016; Pollard et al., 2006). Importantly, such rate analyses can scale in scope, from single positions, to genes, or even discontiguous loci comprising entire pathways (Daane et al., 2020). This contrasts with analyses that quantify a specific measure, like gene loss, expansion, or the detection of identical amino acid substitutions. The latter, for instance, have proven informative in many systems, such as echolocation (Parker et al., 2013), but their significance remains contentious (Marcovitz et al., 2019; Zou and Zhang, 2015). Such specific measures alone fail to capture local, potentially linked, changes that could influence the trait similarly. Single amino acid substitutions are too narrow a measure to encompass the multiple dimensions molecular evolution can act upon. RERs scale to the degree of specificity an experiment requires and can capture disparate yet functionally comparable changes in a single measure.

RERs represent the substitution rate of a single element (exon, gene, CNE, etc.) relative to the genome-wide average. These rates are generally represented with phylogenetic trees and the relative rates can be derived by comparing individual gene trees to the species tree. For example, elephant *PLCZ1* has a much shorter branch length when compared to the average across its genome, resulting in a low RER, while the Elephant Shrew branch is longer than expected and yields a high RER (**Figure 1A**). To identify convergence in rate, the RERs for a specific element can be compared between species and intersected with trait occurrence. Lineages with differing traits and differing RERs are evidence that the locus may be involved in the evolution of that trait. With tens of thousands of loci and only a single species, a distribution of acceleration and constraint would be expected by definition, centered on the background substitution rate. By analyzing many comparisons across multiple instances of trait change, consistently shared RERs for a specific element are unlikely to occur, enabling detection of convergent signals from genomic noise. These probabilities and calculations are also empowered by including the RER magnitude, as they are a continuous variable.

**Figure 1.**
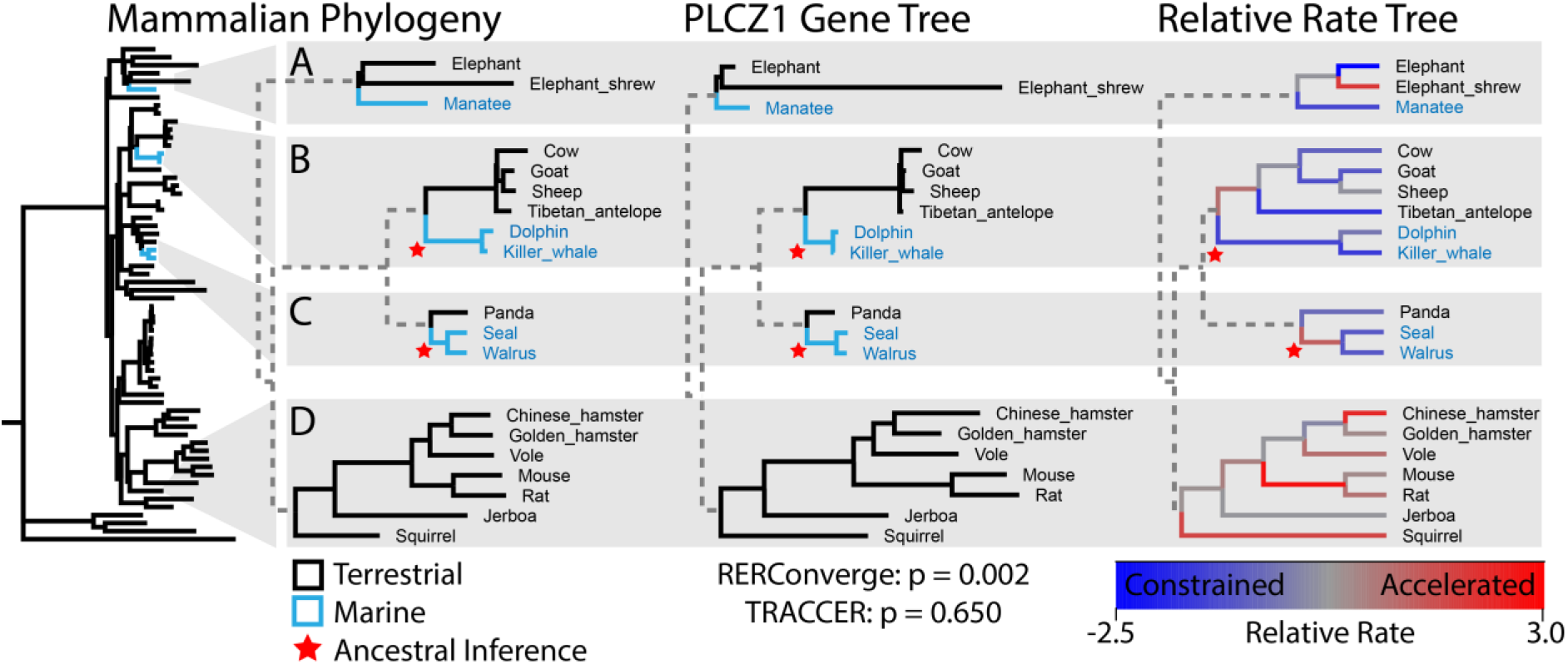
Relative evolutionary rates, topology, and ancestral state assignments in convergence analyses. **A.** Rates of genome-wide variation (the phylogenic tree) are compared to gene-specific rates to calculate RERs. For example, the Elephant and Elephant Shrew terminal branches of gene *PLCZ1* are shorter and longer respectively than their background genomic rates, yielding strong constraint (blue) and acceleration (red) signals on the RER Tree. *PLCZ1* shows constrained evolution on the terminal branches of all five marine mammals (**A,B,C**). Conventional statistics indicates a significant association between *PLCZ1* evolutionary rate and marine transitions (RERConverge p=0.002). However, most of the terrestrial relatives closest to the marine lineages demonstrate similar constraint at this locus (*e.g* elephant, antelope and panda; **A,B,C**, respectively), suggesting *PLCZ1* constraint is a not specific for marine transitions, but a shared property of the broader clades. **C.** Previous analyses delineate the ancestral pinniped as terrestrial, with dramatic influence on the final signal. As 1/7^th^ of the marine branches, its acceleration signal (red) would undermine the convergent constraint shared by the marine terminal branches. Critically, this branch is likely marine, and if either the seal or walrus was missing, the other pinniped would have yielded an acceleration signal for that marine group. **D.** The collection of rodents is consistently accelerated, encompassing a large percentage of the branches on the tree and anchoring the RER distribution, but is distantly related from the marine mammals. By treating every branch as an equivalent and independent instance of marine and terrestrial evolution, conventional methods yield an arguably spurious convergence signal (p=0.002). TRACCER considers that closely related sister lineages are also constrained for this locus, often more so than the marine lineages, while distal clades with no trait change represent much of the acceleration signal. Without needing to infer ancestral states, TRACCER assigns *PLCZ1* a nonsignificant score (p=0.65).

Until recently, tools for RER analysis were limited to individual elements or branches and did not scale to the modern wealth of genomic information. The first broadscale “forward genomics” approach (Hiller et al., 2012) focused solely on trait loss and was agnostic regarding topology. RERConverge later efficiently analyzed RER convergence in acceleration and constraint, implicating genes in marine mammal transitions (Chikina et al., 2016), mammalian subterranean transitions (Partha et al., 2017), and mammalian longevity (Kowalczyk et al., 2020) with fascinating insights into the genetic architecture that facilitates these dramatic environmental and physiological shifts. Those studies assume each branch is an equivalent and independent occurrence of evolution, and as such, process the signals with conventional statistical tests that require independence, including the Pearson Correlation or Wilcoxon Rank-Sum. However, these assumptions are incongruent with the conceptual basis of convergence with consequences for both sensitivity and specificity.

### The Value of Topology

Not all comparisons are equally informative, nor are they truly independent; topology and genetic context must be factored in. Sister lineages with differing traits make for more valuable and informative comparisons than distant ones; the changes facilitating the trait occurred within a shorter-time frame and with a greater degree of shared genetic architecture. As genomic variation must be facilitating the trait change, and there is less variation between sister lineages, any differences revealed are thus more likely to be facilitating the trait difference being investigated. The opposite is equally important; when closely related species with differing traits share a genetic signature, it is strong evidence the signature is unrelated to the trait. Regardless of the arguments dividing convergence and parallelism (Gould et al., 2002; Pearce, 2012), neglecting topology is an oversight of many statistical approaches to associate RERs with trait change. An example of these effects is highlighted in **Figure 1**. There, RERConverge has assigned *PLCZ1* a significant score for convergent constraint in the evolution of marine mammals (**Figure 1A,B,C**). However, the closest terrestrial relatives all share similar levels of constraint, suggesting that constrained PLCZ1 evolution is not specific for marine transitions, but instead a general signature of the larger clades. In contrast, the rodents (**Figure 1D**) are enriched for accelerations and represent a substantial percentage of the branches on the tree, yet are distantly related from any marine lineages. The available mammalian genomes are oversampled for many clades that have little to do with marine transitions. The quantity of branches in these clades can spuriously anchor conventional statistics. Despite these branches representing variable genetic contexts, timeframes, and relationships to the trait being analyzed, conventional statistics must treat them each *as an equivalent and independent instance* of marine or terrestrial evolution. For instance, in the analysis of marine mammals, it is problematic to include the ancestral branch leading to mice and rats -- ̃100M years from the closest marine mammal – as an equivalent comparison and weight as the panda terminal branch, the immediate sister of pinnipeds. Individual branches on a tree are not equally meaningful; they are arbitrary fragments of evolution based on which lineages were sampled, each with their own context and varying degrees of overlap. Instead, the branches from which RERs are derived and compared should represent comparable evolutionary events.

To fully integrate the informative power present in the phylogeny into a convergence analysis requires delineation of which evolutionary events should be compared. Most convergence analyses require flagging specific branches for analysis, with often precarious assumptions of ancestral states. This is easier with well-delineated traits like marine mammals that have a well-documented fossil record (Slater et al., 2012). However, misattribution of these states can easily compromise an analysis. In Chikina et al 2016, the pinniped ancestral branch was not assigned as marine, biasing the constraint signal for *PLCZ1* by leaving out an accelerated branch (**Figure 1C**). If the genome for either the seal or walrus was unavailable, the entire pinniped lineage would have been designated marine and calculated as accelerated. Instead, because the branches are arbitrarily fragmented by sampling availability, both pinniped terminal branches are generating a strong signal of constraint. Critically, these branches likely do not represent a marine transition at all; it is likely the ancestor to the pinniped clade that was undergoing the dramatic adaptation (Berta et al., 2018). By strictly defining the trait status of each branch, there manifests a potential to misattribute when selection may be acting, spuriously raising and lowering signals of selection. This is difficult enough with a clear trait like marine transitions -- the pinniped phylogeny was contentious for some time, and many originally argued it was diphyletic (Koretsky and Barnes, 2003) -- and would be nigh impossible for traits like longevity or metabolism. This problematic requirement is common in convergence analyses. To our knowledge, current rate convergence analyses either ignore topology, or require inferring ancestral states, and/or do not scale to genome-wide data.

## Results

### Topological Weighting and RER Statistics

In light of these concerns, we developed a new analysis pipeline, *Topologically Ranked Analysis of Convergence via Comparative Evolutionary Rates -- TRACCER*. We test if the analysis of convergence is refined by making genomic comparisons between trait-bearing and non-trait-bearing extant species in reference to their most recent common ancestor (MRCA). With this approach, comparisons represent a single genetic context, diverging over the same time frame, and are guaranteed to encompass a change in trait. As each value is now a comparison instead of a singular branch, we can weight each comparison by the phyletic distance between them. This approach allows four critical features: **I**. proximal comparisons can be given greater influence, **II**. phylogenetically distal comparisons can be downplayed, **III**. it is no longer necessary to infer ancestral states, and **IV**. trait distribution can be incorporated to undermine sampling biases.

Once topology is factored in, however, we can no longer use conventional statistics to determine significance as each comparison is no longer truly independent. As we are comparing lineages in reference to their MRCA, many comparisons will share some ancestral branches, causing varying degrees of dependence between comparisons. Instead, we use a permutation strategy to repeatedly shuffle and sample branches of the element trees to simulate how likely a pattern of relative rates is to occur on a given topology with the actual data. The combinatorial complexity of exome-sized datasets on a broadly sampled phylogeny is sufficient to sample from directly. Because RER calculations require a fixed topology, we can shuffle branches between trees to yield a simulated instance of evolution using real examples of how those speciation events occurred at a molecular level. Millions of these phylogenetically and molecularly grounded permutations are analyzed alongside the experimental scores to determine an assumption free background score distribution. The experimental results are compared to this distribution to determine their significance. This contrasts with the variety of evolutionary models one could draw from, which while undeniably useful, their predictions often fail to describe real data (Naser-Khdour et al., 2019). Thus, TRACCER obviates the need for assumptive formulae by drawing directly from the wealth of information inherent to modern comparative genomic datasets.

TRACCER diverges in a few key details of RER calculations from previous analyses with important ramifications. We use a log-transformed fold change from baseline, as opposed to a residual approach used by RERConverge. Residuals fail to inversely allocate weight to variation in longer branches; two SNPs in ten million years should not hold the same influence as two SNPs in one million years, yet these would calculate the same residual value. The fold change approach inherently corrects for branch length and embraces the “rate” concept of relative rates. TRACCER also uses a median distance across all gene trees, excluding zeroes, to derive the relative rates. The median calculation is robust to acceleration outliers, but would be skewed by the number of zero-length branches if they were not excluded. Evolutionary models and tree calculations often struggle with these regions that lack informative substitutions. Due to their short-sequence and/or extreme selective pressure they are guttered to zero-length branches, which does not accurately represent the forces at play genome-wide. These branches should still be analyzed as undergoing constraint, and can still be compared within the same tree, as each branch would be suffering from a consistent bias. The details of these calculations are elaborated in the methods.

These analytical concepts: comparing RERs in reference to MRCAs, weighting comparisons by evolutionary distance, deriving RERs as log-fold changes from zero-corrected-medians, and significance as compared to permutations of the phylogenetic data, empower relative rate analyses to better reveal convergent signatures. TRACCER compares relative evolutionary rates in a pairwise approach that obviates the need for ancestral state assumptions and allows for topologically aware comparisons; it prioritizes sister groups with high comparative power and carefully weighs outgroups to have appropriate influence. RERs are derived as the fold change from the corrected-median branch length, accurately describing rates without branch length bias or assumptive models. Critically, TRACCER defaults include a ranking transformation applied to signal magnitudes, preventing outlier lineages from swamping potential signals, without leaving them ignored. The recurrent concept of convergence is ensured while increasing sensitivity to subtle but meaningful changes over short-time frames. Finally, by scaling each comparison by how often those branches have been used, TRACCER dilutes the influence of over sampled clades and their shared ancestral branches, while allowing their unique terminal branches the full influence. To benchmark the impact of these strategies, we have directly compared TRACCER to a similar analysis, RERConverge, which has recently demonstrated great successes analyzing both marine mammal transitions and mammalian longevity.

### Test Case 1: Marine Mammal Convergence

We obtained amino-acid level gene alignments for 62 mammals from the UCSC 100-Way Vertebrate Multiz dataset (Rosenbloom et al., 2015). We chose this set to match previous analyses with RERConverge (Chikina et al., 2016) and allow direct comparison of statistical resolution. The species tree provided by UCSC was trimmed to maintain just the mammals and used to fix the topology of 37,272 protein coding trees generated with PAML using the same settings as described in RERConverge. The five marine mammals, including the cetaceans (dolphin, killer whale), pinnipeds (seal, walrus) and sirenians (manatee), represent three independent cases of adaptation to an aquatic environment (**Figure 1 Phylogeny**). When using RERConverge, which requires ancestral branches to be classed as well, we additionally flagged the cetacean ancestor as marine, but not the pinniped ancestor, again to match published methodology and results. As a control group, we also copied the published RERConverge selections to allow for direct comparison: aardvark, alpaca, camel, microbat, and myotis bat have roughly similar topology as the marine mammals and provide a non-convergent control group to assess background signal levels.

TRACCER has an integrated diagnostic mode to assess if a particular tree topology has sufficient power to reveal convergence beyond background noise and provide an estimate of how robust of a signal is required to reach that significance. It is important to note that the maximum score for convergence with TRACCER is not derived from merely all trait-bearing species bearing the same extremes of constraint or acceleration; their terrestrial sister lineages must occupy the opposite end of the distribution for that gene. Put another way, sister lineages sharing the same relative rate polarity should compromise the convergent signal, as the signal is no longer specific for the trait in that genetic context. To profile these dependencies, we simulated 10,000,000 gene trees with Brownian motion (Revell et al., 2008) and measured the significance of their convergent signals against a backdrop of another 10,000,000 random trees (**Figure 2A, 0**). This revealed that p-values less than 1/20,000 – roughly the size of a vertebrate exome dataset – were possible, indicating the topology represented by the three marine transitions yields a sufficiently complex score-space to parse significant hits from background noise of a genomic dataset.

**Figure 2:**
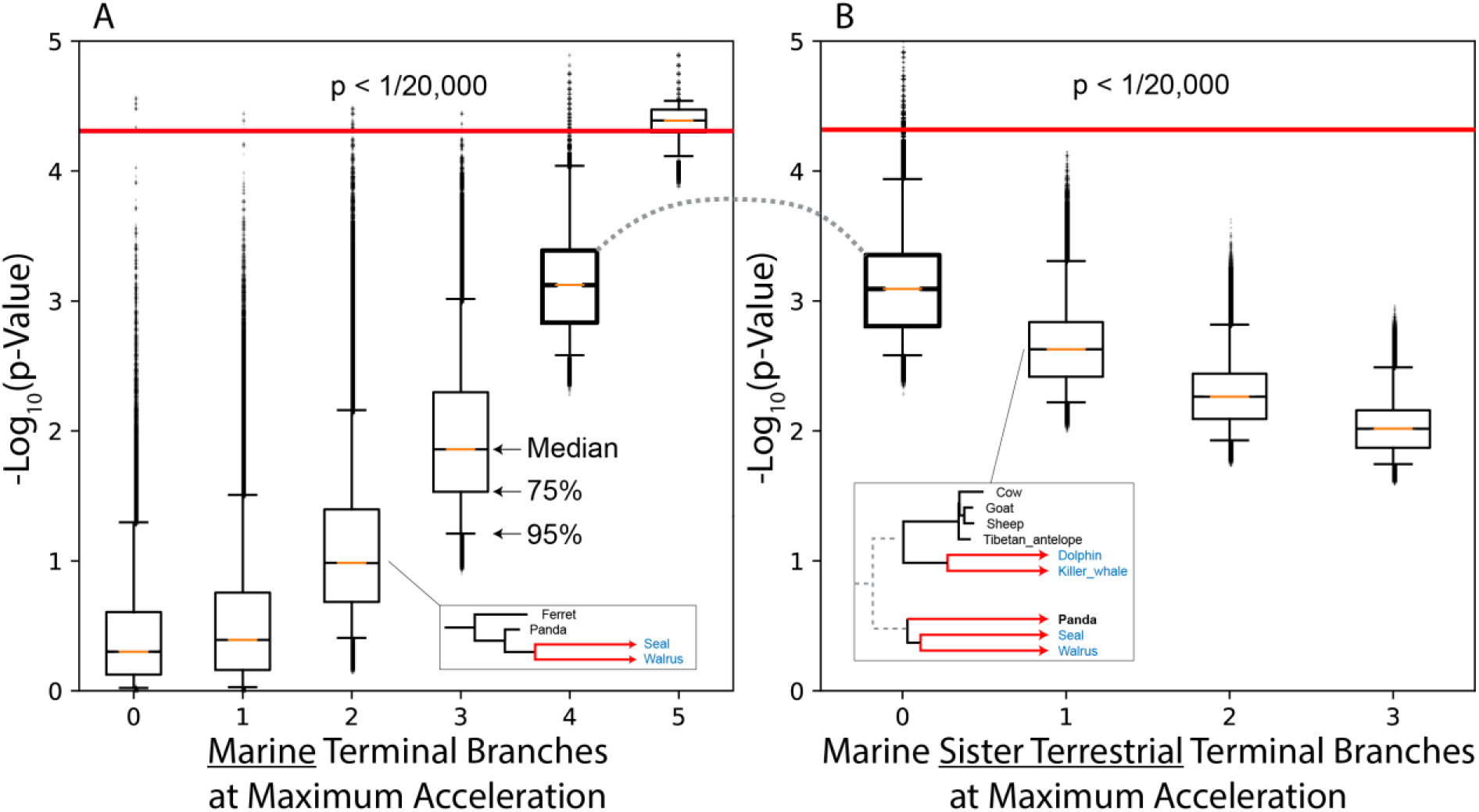
Boundaries of significance for marine mammal convergence. **A.** To highlight how topology influences significance in TRACCER, ten million randomly simulated mammalian gene trees were scored for convergence in marine lineages. Significant probability scores less than 1/20,000 were detected, indicating the topology of the marine dataset is sufficiently complex to identify convergent signals on an exome-sized dataset (**A0, above red line**). To gauge how these p-values were influenced by the pattern of the signal, we forced random marine lineages to extreme acceleration (**A1:5**), with an example of two accelerations in the box. The range of p-values was compressed towards the significance boundary as more lineages were accelerated. However, even with all five accelerated, random variation still produced scores below significance (**A5**). **B.** The source of this variation is highlighted when immediate sister terrestrial lineages are accelerated alongside four marine lineages. While four marine lineages sharing a signal is capable of yielding significant scores, if the closest relative of any marine lineage shares the same extreme acceleration, it is sufficient to prevent those scores from reaching significance. In the example acceleration, the accelerated Panda lineage is undermining the convergent acceleration shared by four random marine lineages.

To roughly gauge the shape or robustness of the convergent pattern required to produce these significant hits, the diagnostic allocates extreme rate accelerations to random terminal branches on top of the Brownian backdrop. We see that with all five marine terminal branches set to maximum acceleration, the vast majority of trees yield a significant score, but not all of them. If the immediate sister lineages were also highly accelerated, they reduce the score below the threshold (**Figure 2A, 5**). We outlined this effect by again purposefully allocating extreme accelerations, this time to four marine mammals along with a number of their immediate sisters (**Figure 2B**). While setting extreme accelerations to four marine terminal branches can drive significant scores, even a single sister lineage sharing that extreme acceleration will entirely prevent it. If the three closest terrestrial lineages share the same extreme signal as four marine mammals, while all other branches are randomly distributed, it will reduce the median p-value by an order of magnitude. These simulations are simplified by distributing the acceleration signals only to terminal branches and randomizing all others but they provide a valuable overview of the topological weight influence.

When applied to the real mammalian genomic dataset, TRACCER reveals substantial enrichment of genes at lower p-values, indicating convergence signals well above the flat background rate seen in the control group (**Figure 3**). In our use, and matching the published methods, RERConverge shows a similar p-value distribution and accurately recapitulates established results with that tool. Unlike those publications, in representing these results and for downstream analyses, we do not segregate accelerated and constrained genes, instead focusing solely on significance. The concordance between these methods is high, with a Spearman correlation of 0.32 and p-value less than 1E-350 (**Figure 4B**). However, a number of genes are dramatically shifted between these analyses, towards both more and less significance. Unlike RERConverge, many TRACCER gene scores remain significant after false discovery correction (**Supplementary S1**) indicating an improvement in sensitivity. To expand upon these differences, GO gene set enrichment was performed using the SUMSTAT approach, which has demonstrated power, flexibility, and simplicity over competing gene set enrichment analyses (Ackermann and Strimmer, 2009; Chen et al., 2010; Tintle et al., 2009). Both TRACCER and RERConverge reveal highly significant processes and systems evolving to facilitate mammalian marine transitions (**Figure 4C, Supplementary S2**). They agree on many of the most significant terms, with extremely significant p-values for cornification, keratinization, myosin filament, and muscle filament sliding. These each have obvious implications with the transition to a marine environment, which required dramatic epithelial adaptions and muscular changes. Another highly significant gene set, “protein-glutamine gamma-glutamyltransferase activity”, is likely playing a role in the marine metabolic and dietary adaptions, particularly with glucose regulation, and has intriguing parallels with diabetes pathology in the literature (Schermerhorn, 2013; Venn-Watson, 2014).

**Figure 3:**
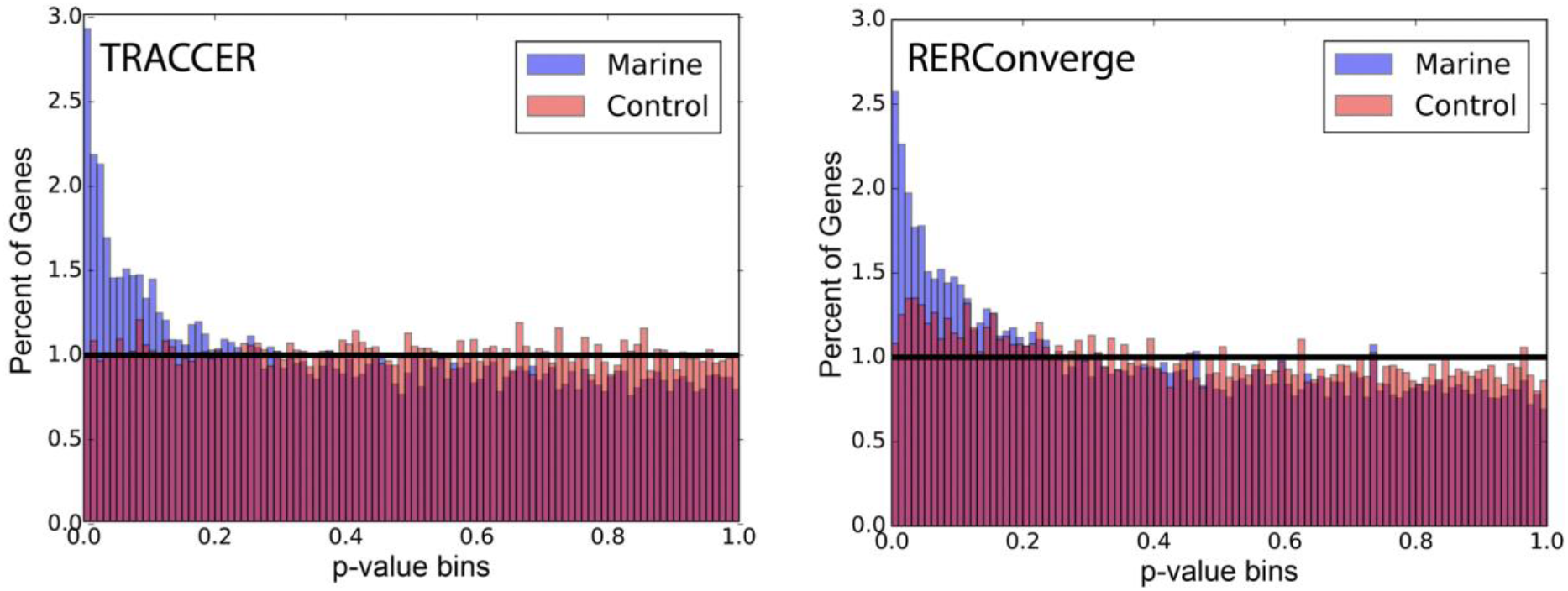
Marine mammal RER convergence significance distributions. Results from both TRACCER and RERConverge analyses are dramatically enriched at low p-values when analyzing for marine convergence (blue). The TRACCER control distribution (red) is flat, matching the distribution expected by chance.

**Figure 4:**
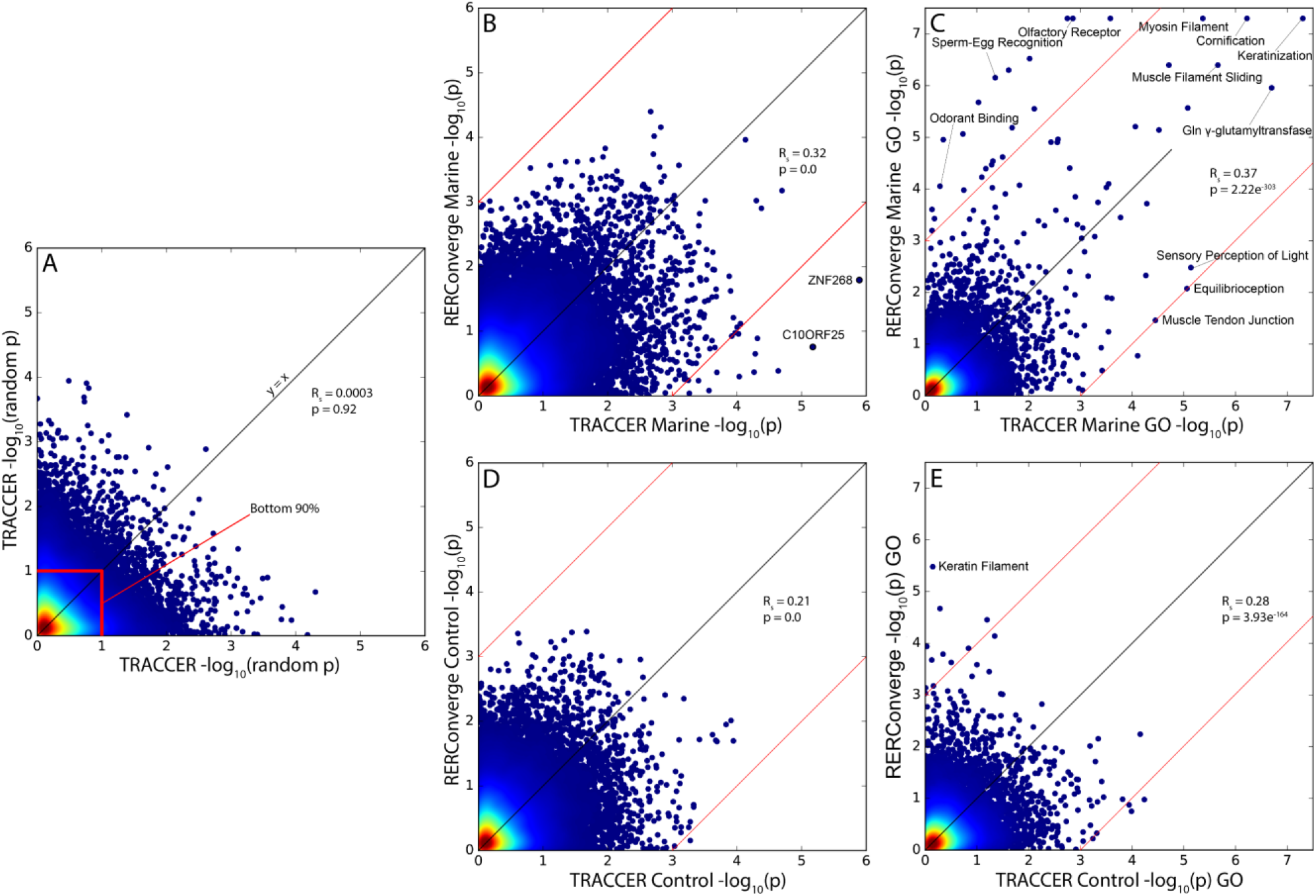
Comparison of TRACCER and RERConverge in marine mammal convergence. **A.** Demonstration of the shape of a random log/log distribution expected by chance with an exome sized dataset using two different, non-convergent species selections. Color indicates density of values, highlighting the shared bottom 90% of p-values concentrated at the origin, and no correlation in results. **B.** Concordance between TRACCER and RERConverge is extremely significant, though TRACCER detects a number of highly significant targets that RERConverge does not (outside of red lines). **C.** GO enrichment of TRACCER and RERConverge gene scores demonstrates agreement on many highly significant terms with established roles in marine transitions (within red lines). However, RERConverge also identifies many terms involved in olfaction and sperm morphology, though there is evidence these are unlikely to be convergently evolving with this definition of marine. **D.** Concordance in the marine negative control dataset is also highly significant, and as expected, neither has any individual genes as highly significant. **E.** GO Enrichment of the negative control dataset, highlighting Keratin Filament as significantly enriched in the RERConverge results, and a lack of false positives with TRACCER.

However, when we focus on the GO-terms with the greatest shift in significance between the two analyses, we reveal some systems are differentially detected, including sperm-egg recognition and odorant binding (**Figure 4C**). Multiple terms involving sperm physiology are differentially detected as significant by RERConverge (**Supplementary S2**). As an example, the ‘sperm-egg recognition’ gene set is one of the most dramatically shifted, with a p-value of 7E-07 using RERConverge, but only 0.044 from TRACCER. We earlier highlighted *PLCZ1* (**Figure 1**) which is the greatest contributor to the ‘sperm-head’ signal in RERConverge. With a p-value shift from 0.002 to 0.65, *PLCZ1* is an excellent example of how topological weighting and ancestral inferences are underscoring the observed differences in the analyses. TRACCER and RERConverge also differ on the conventionally marine-associated GO terms “olfactory receptor activity” and “odorant binding proteins” (ODPs). While there is plenty of evidence that olfaction in general should be reduced, it is not entirely absent, and the discrepancy in these terms may be important. It is certain that many olfactory receptors have been pseudogenized (Hayden et al., 2010), and the olfactory bulb is reduced in most cetaceans and entirely absent in toothed whales (Kishida et al., 2015). Yet, it is only modestly significant in TRACCER (p=.0014) and extremely significant in RERConverge (p < 1E-07). The differential detection of ODPs is driving this shift, as they are entirely a subset of “olfactory receptor activity”, and while significant with RERConverge (p = 8.80E-05), are not with TRACCER (p = 0.52).

The mechanisms and functions of vertebrate ODPs have proven puzzling, but there is evidence they are involved in the perception of pheromones more so than general olfaction (Tegoni et al., 2000). Many marine mammals still utilize pheromones and require the molecular machinery to detect them, so they are unlikely to have been universally compromised as the olfactory receptors have. For instance, dolphins likely employ and detect sex pheromones in their urine (Muraco and Kuczaj, 2015). Critically, the pinnipeds still maintain their vomeronasal system, including strong selective pressure on *TRPC2*, an essential genetic component (Yu et al., 2010), likely because they still maintain some terrestrial behavior. Strictly flagging the pinnipeds as marine mammals, and not their ancestral branch, despite both spending a large fraction of their life on land, may be confounding the analysis. The delineation between terrestrial and marine mammals is not as cut and dry as most convergence publications suggest and echoes the difficulties of inferring ancestral states. The evolution of pheromones in marine mammals and the molecular systems for their detection may be more complex than originally assumed, but according to TRACCER the “odorant binding” gene set is not convergently evolving with this selection of marine mammals, thereby limiting the significance of the more general “olfactory receptor activity” set.

As a negative control for the gene set analysis, we also ran GO enrichment with the same settings on the marine control dataset: aardvark, alpaca, camel, microbat, and myotis bat. TRACCER has no significant terms with FDRs below 0.3 and is within expected levels of background noise. RERConverge, however, has 10 significantly enriched GO-Terms with FDRs below 0.3, including “keratin filament” with a highly significant p-value of 3.3E-06 (**Figure 4E, Supplemental S4**). This raises the concern that the slight enrichment in the control p-value distribution produced by RERConverge (**Figure 2B**) may have a systematic trend; there are no known convergent changes to the integument of the control lineages and there is no immediate explanation for why the keratin gene set should be significantly enriched. In all, TRACCER yielded more significant genes, yet fewer significant GO terms, while maintaining a lower false positive rate in the control GO-enrichment. This suggests improvements in both sensitivity and specificity.

### Test Case 2: Mammalian Longevity Convergence

To assess TRACCER’s ability to parse convergent signals on a more enigmatic, but also more common trait, we applied it to longevity on the same mammalian phylogeny. Again, to compare how the inclusion of topology may augment convergence analyses, we matched the design to previous RERConverge studies (Kowalczyk et al., 2020). As in that study, the longevity trait was calculated as the second principal component of body size and maximum lifespan. These species are effectively “long-lived after correcting for the expected lifespan of their body-size”. However, TRACCER currently only operates on binary traits, so all positive values were treated as equivalently long-lived (**Figure 5A**). We compared these results directly to RERConverge operated with similar settings. Importantly, RERConverge has the functionality to analyze continuous traits; we ran it as binary nonetheless in order to keep the analyses as comparable as possible and gauge the benefit of integrating topology. TRACCER demonstrated substantial enrichment at low p-values, while RERConverge was depleted (**Figure 5B**). The concordance between the studies was significant, with a Spearman correlation 0.12 and p-value 5.53E-120 (**Figure 5C**). Again, many genes were dramatically shifted between the two analytic pipelines, as seen outside the red lines. Importantly, spliceoforms were included, so some proteins are listed multiple times to gain insight into exon specific trends. To generalize the differences observed, we again performed SUMSTAT GO gene set enrichment, using only the most significant of each spliceoform when applicable (**Figure 5D**). TRACCER identifies “Positive Regulation of Growth” (p=5.12E-5), “TORC1” (p = 1.9E-4), and “Telomere Maintenance” (p = 5.96E-5) as three compelling proofs of concept. These terms have obvious implications in the longevity literature (Foley et al., 2018; Papadopoli et al., 2019; Zhu et al., 2019) and lend credence to the others identified in this analysis. Both analyses agree on the significant enrichment of “3’-UTR-mediated mRNA Stabilization”, which has previously been implicated in the evolution of longevity based on patterns of duplication (Doherty and de Magalhães, 2016). However, TRACCER identifies Nucleotide Excision Repair two orders of magnitude more significant than the other terms, and six orders greater than RERConverge scores it. This is driven in part by the most significant member of the group, *POLK*, which has been previously implicated in longevity. Although *POLK* deficient mice are normal in most regards, they do exhibit reduced lifespan (Stancel et al., 2009). More interestingly, POLK is one of the greatest differentially expressed repair genes when comparing human and naked mole rats – relatively long-lived mammals -- to mice (MacRae et al., 2015). The second highest hit in the NER set, *ARL17A*, has its expression significantly associated with human longevity past the 90^th^ percentile (Deelen et al., 2019). NER specifically may be underappreciated in the evolution of longevity.

**Figure 5.**
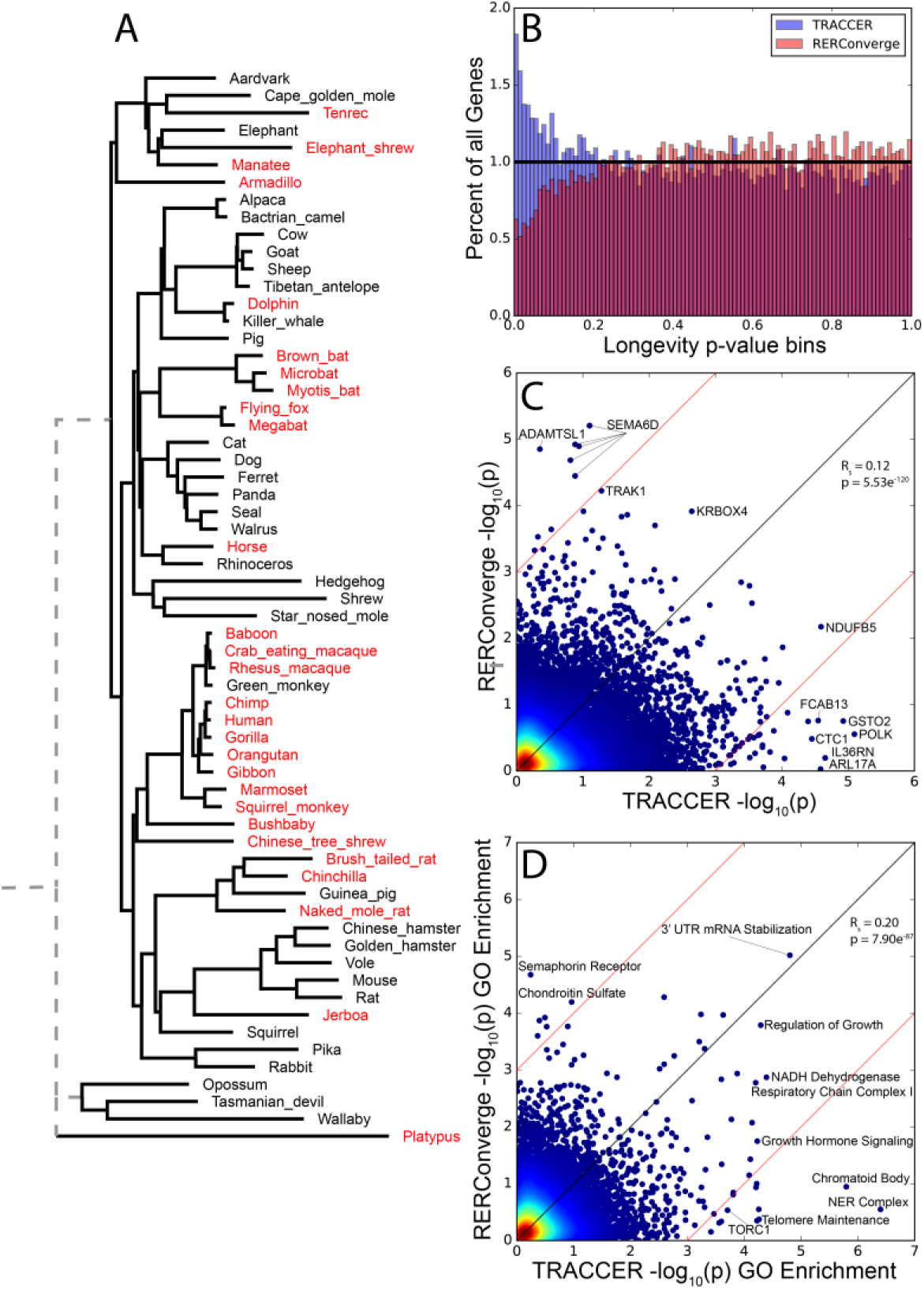
Comparison of TRACCER and RERConverge in convergence of mammalian longevity. **A.** Mammalian phylogeny, highlighting long-lived species after correcting for body size. **B.** Results from TRACCER (blue) are enriched at low p-values when analyzing for longevity convergence, while RERConverge (red) is depleted. **C.** Concordance between TRACCER and RERConverge is significant (p=5.53E-120) with notably different genes labeled. Spliceoforms are included. **D.** Gene set enrichment between the two analyses shows significant concordance, but TRACCER has far more highly significant terms, including proofs of concept like Growth Signaling, TORC, Telomere Maintenance, and Nucleotide Excision Repair. These lend credence to the novel Chromatoid Body set as a promising set of candidates for future longevity investigations. In all, TRACCER yields 20 GO terms convergently evolving with longevity with FDRs below 0.1, while RERConverge has one. Both analyses agree on the significant 3’UTR-mediated mRNA stabilization class.

These proof of concept GO classes lend credence to the highest scoring genes that are not associated with any significant terms and suggests they should be investigated for implications in those systems. *GSTO2, ILL36RN, NDUFB5, EFCAB13*, and *CTC1* all have highly significant scores less than 5E-05 and FDRs less than 0.16, but are not present in the most significant GO terms. They may have unappreciated roles in growth regulation or DNA repair.

The regions of variation and conservation in these genes and gene sets are exciting targets for future longevity investigations.

TRACCER results are also highly enriched for the “chromatoid body” gene set (p=1.10E-6). This functional class of genes is much less studied in longevity than sets like TORC signaling and DNA repair, but has intriguing ramifications and potential. The chromatoid body is essential for maintaining male germ line quality, and its structure and function is demonstrably compromised with age (Santos et al., 2018). Of course, if the male germ line is compromised with age, it would be inherently difficult to select for longevity or the delayed maturation that reliably accompanies it. Many of these genes are validated in germ-line transposable element suppression, but may be more pleiotropic than they first appear, and are potentially involved in a more general transposable element suppression to maintain genomic integrity with age. For instance, the most significant member of this set, *DDX25* (p = 3.7E-4), is an RNA helicase essential for spermatogenesis, but is also broadly expressed in the brain and pituitary where it is poorly characterized. Transposable element suppression may have underappreciated and broad impacts on aging and longevity (Sturm et al., 2015). In all, TRACCER identifies substantially more significant genes and GO terms convergently evolving than comparable methods that remain agnostic to topology (**Supplemental S5,S6**).

## Discussion

Convergence does not occur in isolation; it is inherently a comparative concept, and the various comparisons across a phylogeny are not all equivalent. Comparisons between those with more similar genetic contexts, and importantly, those encompassing trait-changes, will be the most informative to unravel the regulation facilitating that trait change. TRACCER is constructed around this concept in order to augment convergent relative rate analysis with the phylogenetic topological relationships. It builds upon previous work using Relative Evolutionary Rates to infer genomic systems facilitating instances of convergent evolution. TRACCER also includes a diagnostic mode, which is particularly useful for conceptualizing the shape of a convergent signal given a phylogeny of interest. The diagnostic mode is valuable when designing an experiment within a particular group and can guide sampling with an estimation of whether additional species will be worthwhile.

We compare TRACCER directly to RERConverge which uses a similar approach and was the first to use RER convergence across broad datasets (Chikina et al., 2016). We found RERConverge to be powerful and elegant, efficiently revealing significant genes and pathways underlying convergent trait evolution. We use it as a valuable benchmark for TRACCER as it is the most comparable method without factoring in the complexity of topology, though there are other subtle differences in the calculation details. The differences in the results between the two approaches illuminates how the inclusion of phylogenetic relationships and the bypassing of ancestral state inferences may augment convergence analyses. To our knowledge, no other relative rate convergence analysis includes this set of features.

Instead of treating each branch as an independent and equivalent event, TRACCER’s pairwise approach makes comparisons in reference to each pair’s most recent common ancestor. This ensures that each comparison represents the same genetic context evolving over the same time frame, such that the calculated RERs are directly comparable. Each comparison can then be weighted based on the tree topology, without assumptions about ancestral states, and compensates for sampling biases. The default ranking transformations applied to both the RER differences and topological weights maintain the spirit of convergence; it must be a repeated signal, but some comparisons are more informative than others because of their phylogenetic interrelationships. To ignore topology is to leave behind these informative components of phylogenomic data that can refine the map between genotype and phenotype.

RERconverge implicated a number of genes in mammalian marine transitions despite their closest relatives possessing similar signals and/or hinging upon the tenuous but required ancestral state assumptions (**Figure 1**). Part of this is the unfortunate risk of needing to infer ancestral states and flagging the pinniped ancestor as marine would be a simple change that would dramatically change RERConverge results; that branch represents 1/7^th^ of the marine branches in the dataset and can easily shift a signal to and from significance. We repeated this decision when using RERConverge to stay true to published results and to highlight the risks of such requirements. In the case of *PLCZ1*, a gene specific for sperm heads (Escoffier et al., 2016), a significant p-value led to the enrichment of multiple sperm GO terms, none of which have an *a priori* reason to be constrained when adapting to an aquatic environment. To the contrary, scanning electron microscopy available for 36 ungulates and cetaceans suggests sperm-head morphology should be variable in the entire group (Meisner et al., 2005). In fact, the cetacean sperm are missing many of the structures present in the rest of the group, which would not suggest constraint at a genetic level. The loss of such structures is generally associated with a relaxation of selection (Lahti et al., 2009). Inferring ancestral states is often difficult, or even impossible, and discrepancies when assigning them can propagate to databases and downstream functional analyses in the field. Despite these risks, most convergence analyses require it.

By utilizing the full wealth of information available from phylogenomics, TRACCER obviates the need to infer ancestral states and can detect a more refined set of elements than currently available RER approaches. Specificity was improved when applied to the marine mammal dataset. This is particularly evident with the lower false discovery rates, flatter control distribution, and lack of significant hits in the control gene set enrichment as compared to RERConverge. With mammalian longevity, sensitivity was dramatically improved by TRACCER, with many more highly significant genes and gene sets being revealed. These included literature-validated controls like TORC signaling, telomere maintenance, and DNA repair as aspects convergently evolving to modulate mammalian longevity. When robust “proofs of concept” like these emerge, it increases confidence in the novel genes and terms arising alongside, like TRACCER’s highly significant “chromatoid body” GO Term. This gene set is currently only characterized in germ-line transposon suppression, but there is evidence it may be more general in maintaining genomic and epigenomic integrity. The specific highly significant genes that are not members of these defined gene sets are strong candidates for additional investigations and classification. *ZNF268* and *C10ORF25* are the most significant hits for marine transitions but have not been characterized for dermal or muscular functions. *GSTO2, ILL36RN, NDUFB5, EFCAB13*, and *CTC1* are highly significant for convergently evolving with mammalian longevity, but are not present in the most significant GO terms. These results demonstrate that integrating topology into convergent relative rate analyses empowers comparative genomics to discover actionable targets to modulate traits of interest.

## Methods

TRACCER operates on phylogenetic trees, the specifics of which can be decided by experimenter as appropriate for their research. The only requirements are a species tree and a set of individual element trees fixed to the species tree topology. From the species tree, a scalar for each pairwise comparison is derived from the total distance between the two species, with each branch along that path weighted by the number lineages that share it (**Figure 6**). The scalars across the phylogeny are inverted, setting the furthest comparison to the lowest weight, and the closest comparison having the greatest weight. Two options are provided for an additional transformation: the N^th^ root as decided by the user or a rank transformation across all comparisons. The analyses in this study used the rank transformation, with the closest species having the greatest rank. These transformations ensure that proximate comparisons are weighted without swamping the analysis, and distal comparisons are downplayed without being ignored.

**Figure 6.**
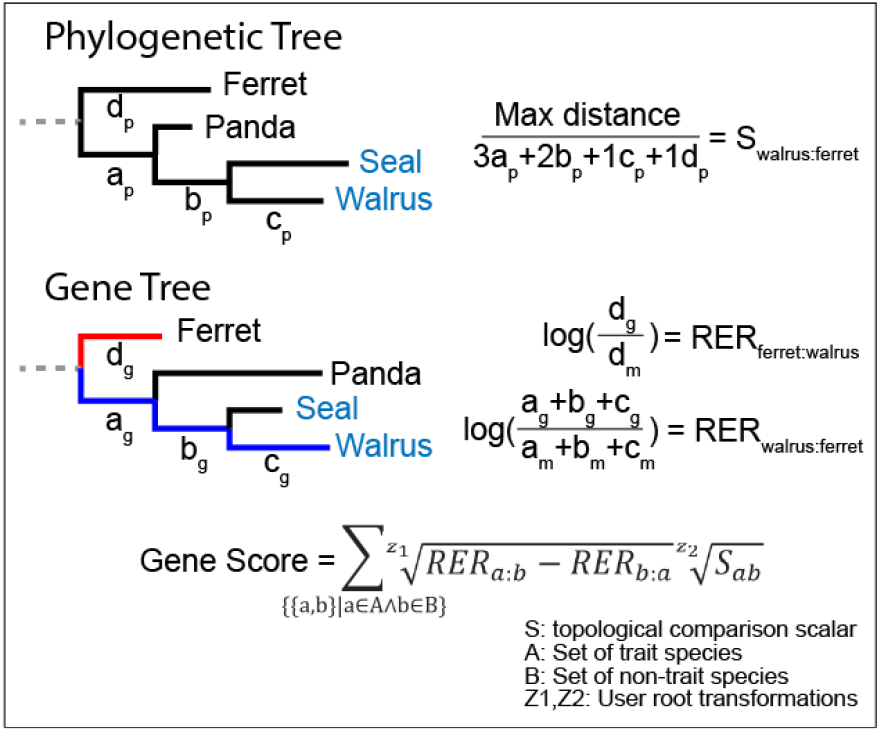
TRACCER Example Calculations. Each pairwise comparison of trait-bearing and non-trait species has a scalar derived from the phylogenetic tree. For example, when comparing Walrus and Ferret, all branch lengths back to their MRCA on the phylogeny (a_p_, b_p_…) are multiplied by the number of extant lineages sharing that branch, and summed. These are then inverted by taking the ratio with the maximum pairwise distance, such that the closest comparison, with the shortest distance between them, has the greatest value. Using the gene tree, RERs for each comparison are derived as the log-ratio of distance to their MRCA (a_g_ + b_g_…) to the median (a_m_ + b_m_…) of that distance across all gene trees. Finally, the difference between RERs for each comparison, and the scalar for that comparison, are z^th^ root transformed as chosen by the user, then summed across all comparisons to produce the gene score. Alternatively, as used in these analyses, they were rank transformed before summing. To calculate significance, these gene scores are then compared to scores calculated from randomly permuted combinations of branches across all gene trees.

The score for a given gene tree is the sum of all pairwise RER differences relative to their MRCA, each scaled by proximity and branch use on the species tree. To calculate RERs for these comparisons, TRACCER uses the medians of the distances across all genes for each lineage to their MRCA, ignoring zeroes and discordance artifacts, to derive a fold-change. Sequence divergence rates are guttered by a large number of zeroes, resulting in a non-normal distribution that makes the concept of ‘relative’ rates meaningless unless they are excluded from this step of the calculation. Importantly, the calculated relative rates are log-transformed. The log transformation balances accelerated and decelerated fold-change relative rates, as a 1/10^th^ deceleration from 1 to 0.1 should be the same importance as the ten-fold acceleration from 1 to 10, despite an order of magnitude difference in the raw differences. As would be expected, the data in our benchmarking investigations is log-normally distributed after removing the zeroes and artifacts.

For each combination of trait- and non-trait bearing species, the log-RERs to their most recent common ancestor are subtracted to quantify differences and direction from their shared ancestral state. By focusing on differences, TRACCER can, for example, detect a relaxation of selection on a conserved element even if the rate is still below the background rate. The pairwise values across the tree are then rank transformed to correct for outliers while ensuring all comparisons have an impact. Then they are multiplied by the scalar derived from the species tree to weigh comparisons by proximity and branch use. All pairwise values are then summed for a final gene score. An example of these calculations is in Figure 6.

To ascertain a given score’s significance, a permutation test is performed to determine the background score distribution. As topology is fixed, branches can be shuffled between trees with millions of permutations and then scored as before. Importantly, when species are missing on a gene tree, it changes the shape of the possible scores and requires its own resampling to compare against. Finally, gene tree scores are ranked against the permuted scores on the same tree topology to determine the percentage that are greater as a p-value. Gene trees exhibiting constraint are only compared to the resamples scoring below the median, and those exhibiting acceleration to resamples above the median.

Fixing trees to the same topology is required to directly compare branches between trees and derive relative rates. This can be problematic when a lineage is missing due to gene loss, as it may remove key branches and undo the 1:1 mapping between trees. Fortunately, because TRACCER performs comparisons in reference to their most recent common ancestor, these can still be compared even when a missing lineage removes an ancestral branch. However, forcing the gene trees to a specific topology will introduce artifacts when the two disagree (Degnan and Rosenberg, 2009) and it would be nonsensical to derive RERs directly from this discordance. Discordance over a larger distance could detected by comparing fixed and unfixed gene trees, or sequence identity directly. Ideally this information would be factored as a complementary dimension of a convergence analysis, but they cannot be used for RERs directly. TRACCER does compensate for one form of discordance: closely related clades and slowly evolving genes often manifest short, but non-zero branch length artifacts.

These artifacts are due to a lack of informative sites in a slowly evolving region. When a lineage has a single SNP while the rest of the clade is identical, the identical lineages will try to cluster. If those are not immediate sisters, it can yield short-non-zero branch lengths. These values can and should be transformed to zero as they represent no change; they are still identical to the ancestral state. TRACCER has an initial curation revealing the distribution of branch lengths to the user, indicating which branch lengths are likely artifacts, and allowing customization of the branch lengths the user wishes to transform. In our test datasets, these took the form of branches with lengths between 10^-4^ and 10^-7^. The exact values will vary based on the tree building software used, but with both PAML and IQ-Tree, in mammals, fish, and birds, we found 10^-4^ to be the right cutoff. While other analyses discard these values, or uses them without correction, TRACCER corrects these values to zero. However, should a gene-tree consist predominantly of these short values, TRACCER will discard it as evolving too slowly to be informative.

Trait distribution and topology is specific to every experiment and dramatically influences the possible score distributions. It may be impossible to differentiate noise from a true convergent signal in a given dataset. To determine if one’s experimental design and topology is sufficient to yield a convergent signal, and how robust that signal must be, TRACCER can be run in diagnostic mode. This feature uses only the species tree to generate millions of gene trees with Brownian motion and forces various convergent patterns upon them. These scores are compared to the strictly random trees to determine how often such scores occur by chance. This can inform experimental design and sampling strategies to optimize resources necessary to get a significant signal.

### Gene Set Enrichment

GO Terms for the mammalian dataset were harvested from Ensembl. Terms with less than three members, or more than five hundred, were discarded as being uninformative or misleading in the context of set enrichment. The SUMSTAT approach was used on log transformed p-values from each analysis, with an additional square root transformation to undermine outliers. In short, the square-rooted, log transformed p-values for each gene in a set were summed and compared to a distribution of randomly sampled scores of the same set size to determine an enrichment p-value. Notably, we do not segregate genes into accelerated and constrained bins for performing enrichment; we use the significance of their convergent signal regardless of the polarity. To handle spliceoforms, only the most significant was included. FDRs are calculated as the expected number of hits at that significance divided by the number of actual hits at that significance or better. At the gene level, the inclusion of spliceoforms makes the calculated FDRs roughly twice as conservative as they should be, but provides potential insight into exon specific influence.

### RERConverge Settings

For marine and control convergence analysis, RERConverge settings were matched to their published protocols (Chikina et al., 2016). For the longevity analysis, the lineages with a positive second principle component of body size and longevity were flagged as bearing the trait (Kowalczyk et al., 2020). These species are effectively “long-lived after correcting for expected lifespan from their body-size”. We used RERConverge’s clade-weighting flag to automatically distribute their weight onto ancestral branches with clade=“all”, weighted=TRUE. When calculating residuals, we used the following settings: transform = “sqrt”, weighted = T, scale = T. Finally, to correlate the residuals with the binary phenotype, we used min.sp=15 min.pos=5, and weighted=“auto”. The minimum species correspond with the minimum species cutoffs used with TRACCER.

### TRACCER as Software

TRACCER’s permutation strategy can be computationally intensive, particularly if variable lineages are missing in many gene trees, as each tree shape will need its own set of permutations to determine its unique score space. However, provided no lineages are missing from any gene trees, or they can be excluded such that only a single score space is needed, TRACCER is efficient and will execute mammalian exome wide datasets in a few minutes. To scale the computational demand for different computing systems, TRACCER can automatically scale the resampling to fit within a designated time window. Alternatively, the user can demand the degree of precision and TRACCER will run until a certain number of resamples are performed. The marine mammal dataset includes 37,272 protein sequences, including spliceoforms, with 6,804 unique topologies. TRACCER runs on this dataset size in 578 minutes with 14 CPUs to achieve a significance limit of 1.6E-7. These times are highly variable depending on the size of the dataset, how many unique topologies there are, and how many unique topologies are significant. The resources required are typically within the scope of any modern computational center, or even a well-equipped personal desktop.

TRACCER was written in Python3 with minimal dependencies for easy installation and execution in a variety of computing environments. Scripts, updates, and tutorials will be made publicly available on github and the lab website at http://www.fishbonelab.org/

